# The rediscovery of *Strix butleri* (Hume, 1878) in Oman and Iran, with molecular resolution of the identity of *Strix omanensis* Robb, van den Berg and Constantine, 2013

**DOI:** 10.1101/025122

**Authors:** Magnus S Robb, George Sangster, Mansour Aliabadian, Arnoud B van den Berg, Mark Constantine, Martin Irestedt, Ali Khani, Seyed Babak Musavi, João M G Nunes, Maïa Sarrouf Willson, Alyn J Walsh

**Affiliations:** The Sound Approach, Rua Dr Pedro Almeida Lima 6, 2710-122 Sintra, Portugal; Department of Bioinformatics and Genetics, Swedish Museum of Natural History, P.O. Box 50007, SE–104 05 Stockholm, Sweden; Department of Zoology, Stockholm University, SE-10691 Stockholm, Sweden; Department of Biology, Faculty of Science, Ferdowsi University of Mashhad, Azadi Square, 9177948974, Mashhad, Iran; Research Department of Zoological Innovations, Institute of Applied Zoology, Faculty of Science, Ferdowsi University of Mashhad, Mashhad, Iran; The Sound Approach, Duinlustparkweg 98, 2082 EG Santpoort-Zuid, Netherlands; The Sound Approach, 12 Market Street, Poole, Dorset BH15 1NF, UK; Khorasan-e Razavi Provincial Office of the Department of the Environment, Mashhad, Iran; Postal code 7661675555, Khucheh Gochin, Golfam Street, Bolvar Shahid Bahonar, Bam, Iran; Parque Ecológico do Funchal, Estrada Regional 103, nr 259, Ribeira das Cales 9050, Monte I Funchal, Portugal; Environment Society of Oman, P.O. Box 3955, P.C. 112, Ruwi, Sultanate of Oman; Wildfowl Reserve, North Slobland, Wexford, Ireland

**Keywords:** molecular identification, nomenclature, phylogenetics, Strigidae, *Strix*, taxonomy

## Abstract

**Background:** Most species of owls (Strigidae) represent cryptic species and their taxonomic study is in flux. In recent years, two new species of owls of the genus *Strix* have been described from the Arabian peninsula by different research teams. It has been suggested that one of these species, *S. omanensis*, is not a valid species but taxonomic comparisons have been hampered by the lack of specimens of *S. omanensis*, and the poor state of the holotype of *S. butleri*.

**Methods:** Here we use new DNA sequence data to clarify the taxonomy and nomenclature of the *S. butleri* complex. We also report the capture of a single *S. butleri* in Mashhad, Iran.

**Results:** A cytochrome b sequence of *S. omanensis* was found to be identical to that of the holotype of *S. butleri*, indicating that the name *S. omanensis* is best regarded as a junior synonym of *S. butleri*. The identity of the *S. butleri* captured in Mashhad, Iran, was confirmed using DNA sequence data. This represents a major (1,400 km) range extension of this species.

**Conclusions:** The population discovered in Oman in 2013 and originally named ‘*S. omanensis’* actually represents the rediscovery of *S. butleri,* which was known from a single specimen and had not been recorded since 1878. The range of *S. butleri* extends into northeast Iran. Our study augments the body of evidence for the recognition of *S. butleri* and *S. hadorami* as separate species and highlights the importance of using multiple evidence to study cryptic owl species.

## INTRODUCTION

Accurate taxonomic designations are important for most, if not all branches in biology. Even in birds, modern scientific studies continue to generate hypotheses of new species, often based on new data and multiple lines of evidence (Sangster 2009, Sangster & Luksenburg 2015). Until the 1960s, studies of the taxonomic status of bird species relied almost exclusively on comparisons of morphological characters. By the 1960s, technological advances made it possible to obtain sound recordings in the field for taxonomic study (Lanyon 1960) and produce audiospectograms (sonagrams) which allowed objective comparison and measurement of acoustic characters. These techniques were first applied to the vocalizations of owls by van der Weyden (1973a, 1973b, 1974, 1975) and Marshall (1978). Subsequent studies of vocalizations have resulted in the discovery of many additional species of owls, a process which continues until the present (e.g. Sangster et al. 2013).

*Strix butleri* was described by Hume (1878) on the basis of a single specimen which was believed to have come from “Omara, on the Mekran Coast” (=Ormara), in what is now southern Pakistan (Fig. 1). Subsequently, small numbers of specimens from Egypt, Israel, Jordan, and Saudi Arabia have been assigned to this species (Goodman & Sabry 1984). In addition, the species is known from Sudan, Yemen and Oman (Mikkola 2012, BirdLife International & NatureServe 2014). However, there have been no subsequent specimens or sight records from north of the Persian Gulf, leading some to suggest that the type of *S. butleri* may have originated from the Arabian peninsula and been brought to Ormara over sea from Arabia (Roselaar & Aliabadian 2009, Kirwan et al. 2015).

**Fig. 1.**
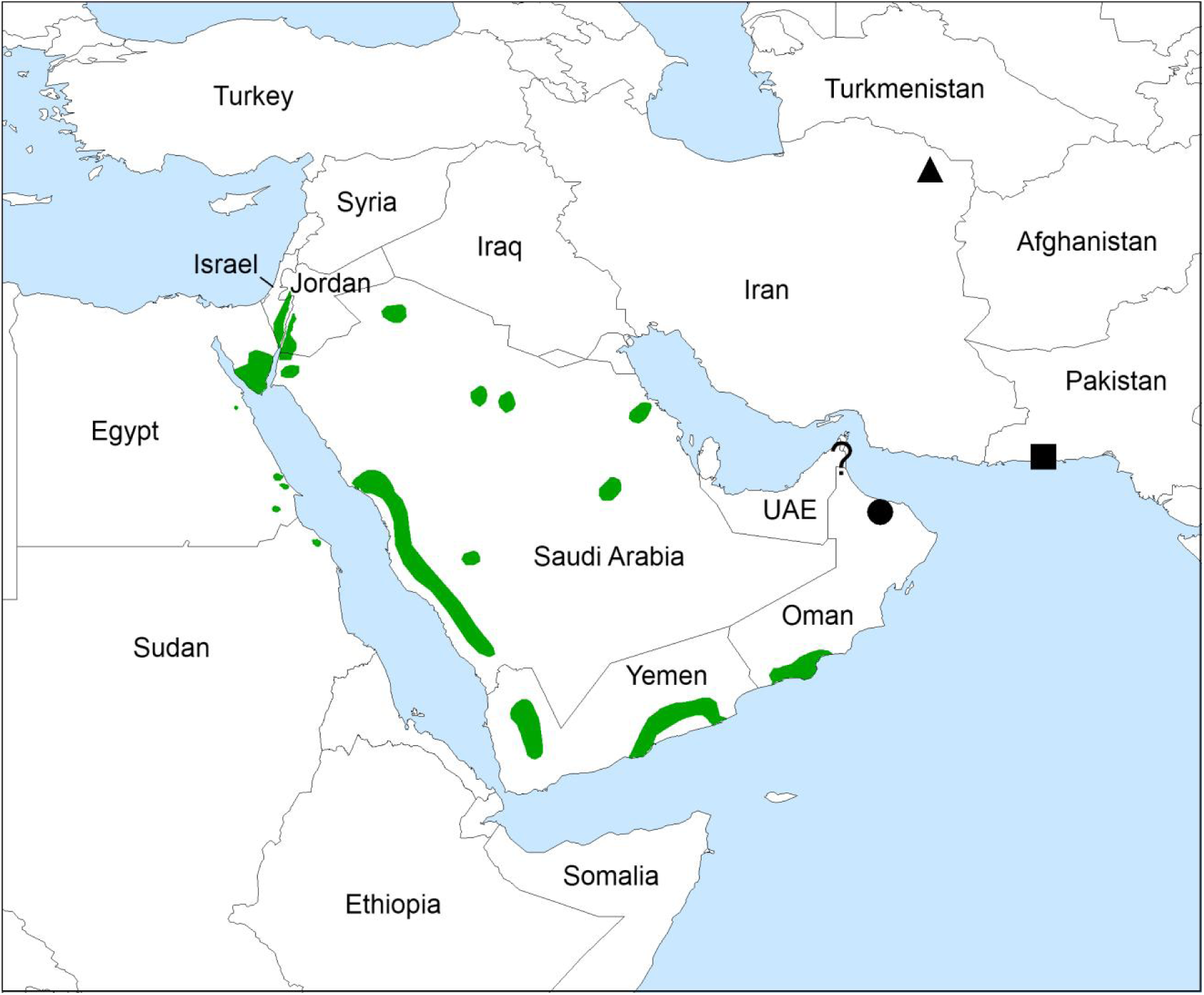
Map showing the known distribution of *Strix hadorami* (green) and *S. butleri* (black). Symbols indicate the type localities of ‘*S. omanensis*’ (circle) and *S. butleri* (square), and the new record in NE Iran (triangle). The question mark denotes a hearing record of *S. butleri* in Wadi Wurayah National Park, United Arab Emirates, which requires substantiation.

In March 2013, Magnus Robb heard vocalisations of an unknown *Strix* owl in the Al Hajar range in northern Oman. In the course of four trips, sound recordings and photographs were obtained demonstrating that the population discovered in Oman represented a different species from ‘Hume’s Owl *S. butleri*’ as it was then understood (Robb et al. 2013). Robb et al. (2013) documented the existence of two species in the Arabian peninsula, based on multiple differences in song, calls, and plumage, and described the Omani population as a new species, *Strix omanensis*. When examining the holotype of *S. butleri* in the Natural History Museum, Tring (BMNH 1886.2.1.994), they did not detect any major differences from the two other specimens of ‘*S. butleri*’ in that collection. Nevertheless, they considered the possibility that the type of *S. butleri* may be same species as *S. omanensis*, and noted that “The eastern location [of the type specimen of *S. butleri*] raises the question whether it in fact could have concerned an Omani Owl [*S. omanensis*]. If it did, the scientific name now used for Hume’s would become the scientific name of Omani while another scientific name would have to be chosen for Hume’s” (Robb et al. 2013).

Kirwan et al. (2015) re-examined the type specimen of *S. butleri* and found that it differed from other specimens attributed to that species in multiple plumage and morphometric characters, indicating that these specimens belong to different species. This was corroborated by analysis of DNA sequences of 218 bp of the mitochondrial cytochrome b gene which showed a sequence divergene of about 10% between the holotype of *S. butleri* and other specimens of ‘*S. butleri*’. They described a new species, *S. hadorami,* to which they assigned all known specimens of ‘*S. butleri*’ except the type of the latter. They did not examine DNA from the Omani population described as ‘*S. omanensis’*. However, they noted that the holotype *S. butleri* showed most of proposed diagnostic character states of *S. omanensis*. Kirwan et al. (2015) suspected that *S. omanensis’* may represent the same species as *S. butleri* and that the holotype of the latter may have originated from Oman.

Critical analysis of type specimens is crucial for the correct application of taxonomic names. Comparisons of the type of *S. butleri* with *S. omanensis* are hampered by the “miserable” state of the former (Meinertzhagen 1930) and the lack of a specimen of the latter. In such cases, comparison of DNA sequences may help to ascertain the taxonomic identity and validity of disputed species-level taxa.

In this study, we use DNA sequences of ‘*S. omanensis*’ to clarify the taxonomic identity of *S. omanensis* and the nomenclature of the *S. butleri* complex. In addition, we use DNA identification techniques to assess the identity of a captured bird (tentatively identified as *S. butleri/S. omanensis*) in Mashhad, Iran, which represents the first record of the species north of the Persian Gulf since 1878.

## METHODS

### Field work: Oman

On 2 March 2015, Alyn Walsh and Magnus Robb caught an Omani Owl at the type locality, Al Jabal Al Akhdar, Al Hajar mountains, Al Batinah, Oman, using a 20 × 4 m mist net. In order to attract an owl to the net, they used playback of several CD tracks from Robb & The Sound Approach (2015) and a decoy owl, painted by Killian Mullarney to look like an Omani and ‘perched’ on a prominent acacia halfway along the net. After catching the owl, they took measurements, feathers, blood samples, photographs and a sound recording. The same measurements were the same as described in Kirwan et al. (2015), taken in the same way. For molecular analysis, they took three feathers from the breast, four tiny ones from the bend of the wing, and two blood samples. In addition they took photographs of the owl in the hand and after release, when it was perched on a thick branch.

The owl was identified as *S. omanensis* (sensu Robb et al. 2013) by the presence of several acoustic and morphological character states which were previously identified as diagnostic for this species (Robb et al. 2013). (i) Shortly before capture, the bird gave diagnostic four-note compound hooting, with the last two notes given in quick succession. In the hand, it showed (ii) orange-yellow eyes, (iii) bicoloured facial disc with dark grey-brown above and beside the eye and pale grey from just above the eye downwards, (iii) very dark, greyish brown upperparts, (iv) ginger-buff to white underparts with long streaks (longitudinal black lines) but only weak transverse bars, and (v) a broad dark trailing edge to the underwing.

### Field work: Iran

In the early morning of 23 January 2015, Ali Khani received news of an owl that had become entangled on the balcony of a house during the night. When he and Babak Musavi went to investigate, they concluded that since it had many feathers of Laughing Dove *Streptopelia senegalensis* around its legs and a blood-covered bill, it may have got in difficulties while hunting. The house was situated in a cultivated area near Vakilabad garden, just west of Mashhad, the second largest city of Iran. South and west of this garden there are barren, rocky slopes possibly offering suitable habitat for Omani Owls. These form part of the northern slopes of the Binalud range, which reaches its highest point (3211 m) at Mount Binalud, some 55 km to the west. Mashhad is c 80 km from the border with Turkmenistan, and over 1300 km from Ormara in Pakistan. They caught the owl, which appeared to be alert and healthy, and collected four feathers for molecular analysis. On releasing it, they took a series of photographs perched and in flight. Having had very little time to prepare for the encounter, they did not attempt to take blood samples or measurements.

### Laboratory procedures and phylogenetic analysis

A blood sample and two feathers from Oman and a single feather from Iran were used for molecular identification. Genomic DNA was extracted using the Qiagen DNeasy Tissue Kit (Qiagen, Valencia, CA) following the protocol of the manufacturer. The lysis procedure was prolonged to 18 hours, and 20 μl of 1 M dithiothreitol (DTT) solution was added during to the initial lysis step.

The mitochondrial cytochrome b (cyt b) was amplified because this is the only marker for which sequences of the holotypes of *S. butleri* (BMNH 1886.2.1.994) and *S. hadorami* (BMNH 1965.M.5235) are available (Kirwan et al. 2015). Amplification was performed in two overlapping fragments. Primer sequences were newly designed, and are as follows: CytbStrixF1 (5’-GAATCTGCCTAATAGCCCAAATC-3’), CytbStrixR2 (5’-AAGCCACCTCAGGCTCATTCTAC-3’), CytbStrixR3 (5’-GGAGAGTGGGCGAAAGGTTATT-3’). The primer combination F1/R2 amplifies 345 bp and F1/R3 amplifies 806 bp. Both fragments fully cover the sequences of the holotypes of *S. butleri* and *S. hadorami*.

PCR products were cycle-sequenced in both directions using the Big Dye Terminator v1.1. Sequences were read on an ABI 3100 capillary sequencer (Applied Biosystems, Foster City, CA). Sequence fragments were aligned and visually edited using Lasergene Editseq (DNA Star, Madison, WI). Both sequences are deposited at GenBank (accession numbers KT428757–KT428758). DNA sequences of six other species of *Strix* were obtained from GenBank. *Tyto alba* was used as an outgroup. Genbank accession numbers and references to the original sources are given in Table 1.

**Table 1.** Genbank accession numbers of samples used in molecular analyses.

| **Taxon** | **GenBank accession number** | **Source** |
| --- | --- | --- |
| *Strix omanensis* (Oman) | KT428757 | This study |
| *Strix butleri* (Iran) | KT428758 | This study |
| *Strix butleri* (holotype) | KM459027 | Kirwan et al. (2015) |
| *Strix hadorami* | AJ003912 | Wink & Heidrich (1999) |
| *Strix hadorami* | AJ003913 | Wink & Heidrich (1999) |
| *Strix hadorami* | EU348994 | Wink et al. (2009) |
| *Strix hadorami* (holotype) | KM459028 | Kirwan et al. (2015) |
| *Strix woodfordii nigricantior* | EU348995 | Wink et al. (2009) |
| *Strix woodfordii* | AJ004065 | Wink & Heidrich (1999) |
| *Strix woodfordii* | AJ004066 | Wink & Heidrich (1999) |
| *Strix woodfordii woodfordii* | AJ004064 | Wink & Heidrich (1999) |
| *Strix uralensis* | JX092123 | Hausknecht et al. (2014) |
| *Strix uralensis* | AB741546 | Omote et al. (2013) |
| *Strix aluco* | AJ004045 | Wink & Heidrich (1999) |
| *Strix aluco* | AJ004057 | Wink & Heidrich (1999) |
| *Strix nebulosa* | AJ004058 | Wink & Heidrich (1999) |
| *Strix nebulosa* | AJ004059 | Wink & Heidrich (1999) |
| *Strix rufipes* | AJ004060 | Wink & Heidrich (1999) |
| *Strix rufipes* | AJ004061 | Wink & Heidrich (1999) |
| *Strix varia* | AF448260 | Desmond et al. (2001) |
| *Tyto alba* | FJ588458 | Braun & Huddleston (2009) |

Phylogenetic relationships were estimated with maximum likelihood (ML) analysis using MEGA5 (Tamura *et al*. 2011). Clade support for the ML analysis was assessed by 1000 bootstrap replicates. The best-fit model was estimated with MEGA5 using the Akaike Information Criterion. The selected model was HKY + G. To further evaluate statistical support for the topology, we ran a Bayesian analysis using MrBAYES version 3.2.2 (Ronquist et al. 2012). Default priors in MrBAYES were used. We ran four Metropolis-coupled MCMC chains for 1 million generations and sampled the topology every 100 generations. Convergence between the two MrBayes runs was assessed by comparing the posterior probability estimates for both analyses using the program AWTY (Nylander *et al*. 2008). The first 25% of the generations were discarded (‘burn-in’) and the posterior probability was estimated for the remaining sampled generations. Uncorrected p pairwise sequence divergences were calculated in MEGA5 with complete deletion of nucleotide positions with missing data.

Nuclear copies of mitochondrial sequences (numts) may represent a problem in mtDNA studies (e.g. Den Tex *et al*. 2010). We used several lines of evidence to assess the authenticity of our sequences. First, electropherograms were inspected for double signal (two clear peaks at one or more nucleotides), which indicates a mixture of mitochondrial and nuclear sequences (Den Tex *et al*. 2010). Second, we checked the translated consensus sequence for the presence of frameshift mutations or stop codons, which are strong indications that a sequence does not represent that of a protein-coding gene. Finally, we checked whether nucleotide substitutions were primarily found at the third codon, which is expected when a sequence is of a protein-coding gene. In old numts, the distribution of substitutions is expected to be equal across all three codon positions (Zink & Barrowclough 2008).

## RESULTS

### Morphology: Oman (Fig. 2a and Fig. 2b)

**Fig. 2.**
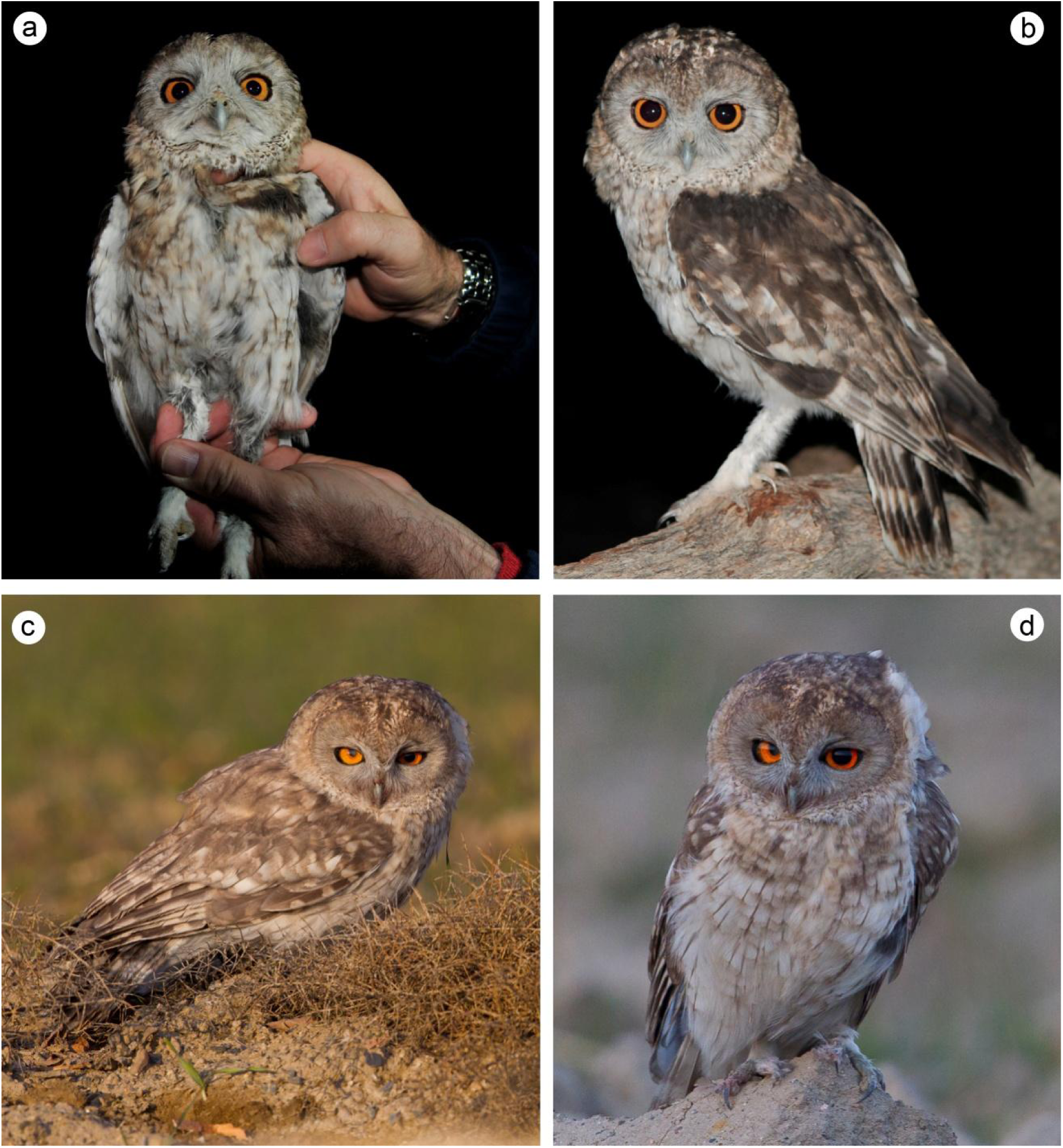
Photographs of (a, b) *Strix butleri* captured at the type locality of ‘*Strix omanensis*’, Al Hajar range, Oman, 2 March 2015 (Magnus S. Robb & Alyn J. Walsh) and (c, d) *Strix butleri* after release, Mashhad, Iran, 23 January 2015 (Seyed Babak Musavi).

Morphometric data of the captured bird are given in Table 2.

**Table 2.** Morphometric data obtained from an individual of ‘*S. omanensis*’ (=*S. butleri*) caught in the Al Hajar range, northern Oman on 2 March 2015.

| Variable | State |
| --- | --- |
| Tarsus | 67.4mm |
| Wing | 255 mm |
| Tail | 142 mm |
| Tail graduation | 15 mm |
| Bill (upper mandible from skull to tip) | 31.85 mm |
| Bill (skull to nostrils) | 17.7 mm |
| Bill (skull to centre of curve) | 24 mm |
| Bill depth at end of feathering | 14.0 mm |
| Bill depth from top of cere | 16.0 mm |
| Weight | 220 g |
| Moult | p1 + p2 old on left wing |
| Primary 1 to wingtip | 56 mm |
| P2 to wingtip | 13 mm |
| P3 to wingtip | 0 mm |
| P4 to wingtip | 0 mm |
| P5 to wingtip | 8 mm |
| P6 to wingtip | 33 mm |
| P7 to wingtip | 50 mm |
| P8 to wingtip | 60 mm |
| P9 to wingtip | 71 mm |
| P10 to wingtip | 80 mm |
| Secondary 1 – wingtip | 93 mm |
| P1 falls | between 7 + 8 |
| P2 falls | between 5 + 6 |

#### Structure

Medium-sized owl with rounded head lacking ear-tufts, a well defined facial disc and typically large eyes. Tarsi long. Tail short. Wing-tips level with, or projecting marginally beyond end of tail, depending on posture.

#### Head

Facial disc pale grey, gradually becoming darker grey-brown above eye. Upper half of disc narrowly bordered dark brown; lower half with creamy or light buff ‘ruff’, finely stippled with dark spots. Prominent dark median crown-stripe beginning just above eye level, widening slightly toward top of head and contrasting with two narrow clusters of whitish-tipped feathers either side, running from forehead onto crown. Pale grey forward-pointing facial feathering just above eye and bristly ‘moustache’ hardly contrasting with lower half of facial disc. Crown densely mottled dark on a lighter ground, sides of head with more ginger ground colour, gradually shading to off white toward lower nape. All feathers of sides and back of head pale-based and dark-tipped resulting in irregular pattern of light spots and dark blotches or bars following the contours of feather tracts. Largest whitish spots concentrated in nuchal band at back of head. Chin whitish, throat light buff, finely stippled dark.

#### Upperparts

Mantle, scapulars, back, rump and uppertail-coverts dark grey-brown with diffuse buff and whitish spots of varying size and intensity.

#### Underparts

Breast washed light ginger-buff, strongest (verging on rust-coloured) at sides, with loose arrangement of narrow dark shaft-streaks and few faint transverse bars. Belly and flank whitish with longer thin shaft-streaks and sparsely distributed, faintly marked buff-brown bars. Abdomen, undertail-coverts and thigh off-white, unmarked.

#### Upperwing

Primaries barred dark brown and greyish-buff, five light bars (including tip) interspaced with four broader dark bars. Secondaries similar but fewer bars (three light, three dark) and pattern with slightly less contrast than on primaries, especially toward base. Tertials brown, innermost with three narrow but distinct buff bars on the inner web, the middle and subterminal bars continuing onto the outer web. Alula dark grey-brown, longest feather apparently fresher and with three buff notches on outer web, shorter feathers plain. Greater and median secondary coverts brown with large whitish subterminal spot on outer webs of outermost feathers, smaller and less distinct pale markings on coverts closer to body. Lesser and marginal coverts more uniform dark brown. Greater primary coverts almost uniform dark brown with very subdued barred pattern.

#### Underwing

Outermost primary plain brown-grey with faint longitudinal streak on middle of inner web, rest of primaries boldly barred brown and white/buff-grey, contrast between light and dark bars more pronounced at base where, toward inner primaries, white bars broadened and proximal dark bar much reduced in strength. Secondaries similar to inner primaries, extensively white at base merging imperceptibly with clean white greater coverts. Greater primary coverts white with bold dark tips to outer six feathers forming a prominent dark carpal-crescent. Remaining underwing coverts greyish with fine dark shaft-streaks, marginal coverts (leading edge of wing) white.

#### Tail

Upperside boldly barred dark brown and greyish-buff, three broad dark bars, and three or four narrow light bars, including tip. Light bars on central pair of rectrices reduced, especially on inner webs, so these feathers darker and less strongly patterned than the rest. Underside similarly marked to uppertail but pattern even bolder due to light bars being almost whitish. Three dark bars and up to three light bars visible beyond undertail coverts, width of light and dark bars more equal than on upperside.

#### Bare parts

Pupils black, iris orange-yellow with black surround; eyelid dark greyish. Bill pale green-grey. Tibia, tarsus and toes feathered whitish, soles light yellowish-buff, claws light horn-grey.

### Morphology: Iran (Fig. 2c and Fig. 2d)

#### Structure

Medium-sized owl with rounded head lacking ear-tufts, a well defined facial disc and typically large eyes. Tarsi long. Tail short. Wing-tips level with, or projecting marginally beyond end of tail, depending on posture. Possibly not as long-legged as Omani individual; this may simply be due to the bird having been photographed in a more relaxed stance, with body plumage fluffed out concealing the true length of the tarsus.

#### Plumage, general

Overall impression is of bird that is lighter in colour, especially on the upperparts and folded upperwing, than individual from Oman. However, since all existing photos of ‘*omanensis*’ have been taken either at night, using flash, or of birds sitting within roost-holes by day, comparisons with photos of Iranian owl (in low evening light, without the use of flash) need to be made with caution.

#### Head

Very similar to captured Omani individual. Buff colour on sides of head bordering upper part of facial disc a little paler and more washed-out but this is of doubtful significance. Facial disc grey, gradually becoming darker grey-brown above eye. Upper half of disc narrowly bordered dark brown; lower half with creamy or light buff ‘ruff’, finely stippled with dark spots. Prominent dark median crown-stripe beginning just above eye level, widening slightly toward top of head and contrasting with two narrow clusters of whitish-tipped feathers either side, running from forehead onto crown. Pale grey forward-pointing facial feathering just above eye and bristly ‘moustache’ hardly contrasting with lower half of facial disc. Crown densely mottled dark on a lighter ground, sides of head with paler buff ground colour, gradually shading to off white toward lower nape. Chin whitish, throat light buff, finely stippled dark.

#### Upperparts

Mantle, back, rump and upper-tail-coverts not visible in photographs; scapulars with buff and whitish spots but apparently lighter grey-brown ground colour than in captured *‘omanensis’*. Note, however, that in one photo (Fig. 2d) where bird not illuminated by sun, brown of the upperparts and head appears considerably darker in tone.

#### Underparts

Breast washed light apricot-buff, strongest at sides and extending further down towards legs than in captured *‘omanensis’*, with loose arrangement of narrow dark shaft-streaks and few faint transverse bars. Belly, flank and undertail coverts whitish with longer thin shaft-streaks and sparsely distributed, faintly marked buff-brown bars. Abdomen and thigh off-white, unmarked.

#### Upperwing

Mostly based on photos of folded wing, though unsharp flight photo also informative. Remiges barred dark brown and pale buff, with pale buff tip. Tertials not clearly visible in photos. Alula dark grey-brown, all feathers notched with buff on outer web. Greater and median secondary coverts fairly pale brown with large whitish subterminal spot on outer webs of outermost feathers, smaller and less distinct pale markings on coverts closer to body. Lesser and marginal coverts more uniform brown. Primary coverts distinctly barred, much more so than in captured *‘omanensis’*.

#### Underwing

Not visible in photos.

#### Tail

Only partly visible in sharp photos, though upperside visible in unsharp flight photos. Upperside boldly barred dark brown and pale buff, three broad dark bars, and four narrow light bars, including tip. Underside similarly marked to uppertail but width of light and dark bars more equal. Three dark bars and up to three light bars visible beyond undertail coverts.

#### Bare parts

Pupils black, iris orange-yellow with black surround; eyelid dark greyish. Bill pale green-grey. Tibia, tarsus and toes feathered whitish, soles light yellowish-buff, claws apparently a bit blacker than in captured *‘omanensis’*, but probably due at least in part to different light conditions.

### Molecular identification

We obtained 790 base pairs (bp) of cytochrome b of *S. omanensis* and 767 bp from the owl caught at Mashhad, Iran. We found no evidence of numts. Electropherograms showed no double signal; the alignment showed no stop codons, insertions or deletions; and most (65/78, 83%) nucleotide substitutions relative to the longest *S. hadorami* sequence available on GenBank (EU348994) were found in the third codon and resulted in only three amino acid substitutions.

The sequence of *S. omanensis* was identical to the short (218 bp) sequence available from the holotype of *S. butleri* (Genbank acc. no. KM459027). The sequences of *S. omanensis* and the Iranian owl were almost identical, differing in only two nucleotides (0.26%), both at third positions. Across 790 shared bp, the sequence of *S. omanensis* differed from that of *S. hadorami* (EU348994) by 78 substitutions, corresponding to an uncorrected sequence divergence of 9.9%.

Phylogenies based on ML and BI produced identical phylogenies in which both *S. omanensis* and the owl caught at Mashhad, Iran clustered with the holotype of *S. butleri* (Fig. 3). This was strongly supported in both ML (98%) and Bayesian analyses (1.0 PP). In these analyses, *S. hadorami* and *S. butleri* formed reciprocally monophyletic groups. Relationships with *S. woodfordii* were unresolved, most likely due to the small number of nucleotide sites analysed.

**Fig. 3.**
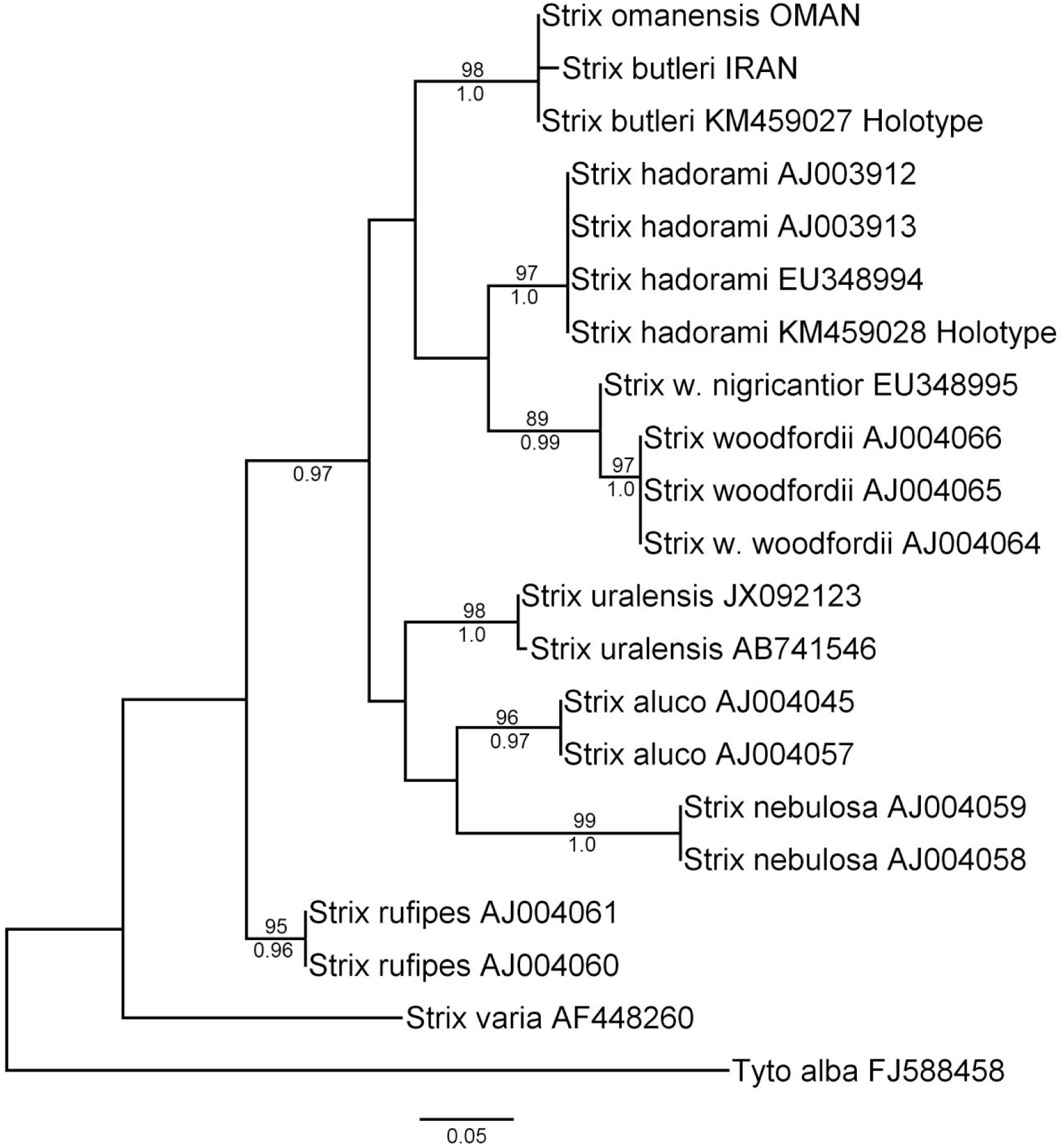
Maximum likelihood phylogeny of *Strix* owls based on 218 bp of cytochrome b, showing the position of *Strix omanensis* Robb, van den Berg & Constantine, 2013 sampled at its type locality and the owl sampled in Mashhad, Iran in January 2015. Maximum Likelihood bootstrap support values (>80%) and Bayesian Posterior Probabilities (>0.95) are given above and below branches, respectively.

## DISCUSSION

### Taxonomy and nomenclature

Mitochondrial DNA (mtDNA) has long been a popular marker in taxonomic and molecular identification (‘barcoding’) studies of birds. This is due to its presence in high concentrations in tissue material, its smaller effective population size which results in faster fixation rates compared to nuclear DNA and, as a consequence, its ability to distinguish a large proportion of species (Zink & Barrowclough 2008, Ward 2009). Our study found that the cytochrome b sequence of a member of the population described as *S. omanensis* (Robb et al. 2013) and sampled at its type locality is identical to that of the holotype of *S. butleri*. This is a strong indication that *S. omanensis* and *S. butleri* belong to the same evolutionary lineage. However, there are some examples of valid species of birds that cannot be reliably distinguished using mtDNA markers. In most of these there is strong evidence from other data that these represent species (e.g. Crochet et al. 2002, Joseph et al. 2006, Irwin et al. 2009, Joseph et al. 2009, Campagna et al. 2010, Päckert et al. 2012). Thus, a lack of fixed mtDNA differences cannot by itself be considered falsification of the existence of species taxa (de Queiroz 2007). Despite this caveat, we believe that current evidence does not justify maintaining *S. omanensis* as a separate species because there is no positive evidence that it represents a separate lineage from *S. butleri*. Therefore, the name *Strix omanensis* Robb, van den Berg and Constantine, 2013 is best treated as a junior synonym of *Asio butleri* Hume, 1878 (now *Strix butleri*).

By providing evidence that the population in Oman previously known as ‘*S. omanensis*’ is *S. butleri*, our study augments the body of evidence supporting the treatment of *S. butleri* and *S. hadorami* as separate species. Whereas the evidence available to Kirwan et al. (2015) was limited to a specimen of *S. butleri* and two lines of evidence (DNA and morphology) differentiating it from *S. hadorami*, the hypothesis that these are species is now also supported by bioacoustic evidence, plumage data from photographs of multiple individuals of *S. butleri*, and DNA sequences of three individuals.

Demographic and genetic exchange between Omani and Iranian populations of *S. butleri* is probably limited by the Gulf of Oman and the Strait of Hormuz. Future studies should focus on making objective comparisons of the plumage and vocalizations of Omani and Iranian populations of *S. butleri*. This is not currently possible due to the absence of specimens from both countries, and of recordings from Iran, where there have been no further observations. More detailed molecular comparisons are warranted to investigate possible population structure and genetic diversity within *S. butleri*, which could inform both taxonomic and conservation genetic studies.

To avoid confusion, we propose to exclude ‘Hume’s Owl’ (and ‘Hume’s Tawny Owl’) as the English name for either species because this is an ambiguous name. Until the end of 2014, it was used universally for what is now *S. hadorami*. At the same time it has historical links to *S. butleri*, the species actually described by Hume. Retaining it for either species may result in misunderstanding. Kirwan et al. (2015) proposed the name ‘Desert Tawny Owl’ for *hadorami*, but this may be shortened to ‘Desert Owl’ to avoid the implication of a close relationship with Tawny Owl *S. aluco* or having to add a modifier such as ‘Forest’ to the latter name. We recommend the name ‘Omani Owl’ for *S. butleri* sensu stricto, because the only known population of this species is in Oman, with only single individuals ever having been located outside Oman.

### Rediscovery and distribution of *S. butleri*

Our study documents the extension of the range of *S. butleri* by 1,300 km to the Mashhad region in northeastern Iran, and its presence in the Al Hajar range of northern Oman (Fig. 1). Its range in Arabia may extend west to Wadi Wurayah National Park in the United Arab Emirates where it was identified in March 2015 by vocalizations (Jacky Judas pers comm) although further substantiation is desirable. Clearly, *S. butleri* is a highly elusive species which is difficult to study in the field. Further field work in Oman, the United Arab Emirates, Iran and Pakistan, perhaps aided by the use of song playback, is necessary to elucidate the range of *S. butleri.*

## ACKNOWLEDGEMENTS

This study was financed and supported by The Sound Approach. It forms part of a broader Omani Owl conservation project conducted as a collaboration between The Sound Approach, BirdLife International and the Environment Society of Oman. We would like to thank the Omani Ministry of Social Development for approving this collaboration, and the Ministry of Environment and Climatic Affairs for granting us permission for fieldwork and to take genetic samples from a wild Omani Owl (permit nr 5/2015). The Office for Conservation of the Environment also advised us during the fieldwork phase of the project. We would like to thank Parque Ecológico do Funchal for making it possible for João Nunes to join us, and David Tierney for his forbearance with Alyn Walsh’s long absences. Andrew Spalton provided welcome advice and good company in the field. In the days immediately following the discovery in Iran, Richard Porter arranged for The Sound Approach and the Iranian team to work together, for which we are extremely grateful. We thank Niloofar Alaye for her assistance with the molecular work. Killian Mullarney gave valuable feedback on the manuscript.

